# Monoallelic expression characterizes a distinct molecular and clinical group of breast tumors

**DOI:** 10.64898/2025.12.09.692870

**Authors:** Mona Arabzadeh, Amartya Singh, Kyle Payne, Shridar Ganesan, Hossein Khiabanian

## Abstract

In diploid cells, allelic imbalance occurs when gene alleles are expressed at different levels. To investigate the allelic imbalance landscape in tumor samples, we developed Interval-Based Allelic Imbalance Detection (IB-Aid), a quantitative framework that uses interval arithmetic to robustly distinguish monoallelic from biallelic gene expression by computing confidence intervals based on sequencing measurement uncertainty. We applied this approach to The Cancer Genome Atlas Breast Invasive Carcinoma tumor samples and through unsupervised gene enrichment analyses, identified a group of patients with a distinct monoallelic gene expression signature. Notably, these patients were enriched in Black/African American patients and had tumors which had not been previously classified into any established molecular subtype. Clinically, these tumors were associated with poor overall survival, with survival outcomes comparable to the aggressive basal subtype. These findings suggest a potential link between allelic imbalance and breast cancer development and point to genetic and epigenetic mechanisms that drive allelic imbalance as novel biomarkers for prognosis and design of targeted treatment strategies.

## INTRODUCTION

In a diploid genome, the relative abundance of transcripts that express the coding sequences in either copy of the genome is modulated by nearby cis-regulatory promoters, enhancers, and silencers in the non-coding regions of the genome that control the expression of nearby genes as well as by trans-regulatory factors that modify the expression of distant genes through three-dimensional interactions. From an allele-specific point of view, cis-acting factors affect expression locally for either the maternal or the paternal alleles while trans-factors may affect both alleles^1^. Potential germline variants have been shown to be associated with unequal, imbalanced expression of the alleles^2–4^. In addition, 1-15% of autosomal genes may show random allelic expression^5^, associated with rapidly evolving regions in the human genome, adaptive signaling processes, and genes linked to age-related diseases such as cancer^6^; however, establishment of causal relationships between allelic DNA copy and allelic gene expression requires deeper functional inquiry.

Random monoallelic expression (RMAE)^7^ and X-chromosome inactivation^8^ are common in normal tissues; however, only clonally related cells arising from the expansion of a cell with a specific RMAE pattern will share the same allelic expression profile. Consequently, bulk transcriptomic measurements of relative allelic expression typically average to a balanced profile across heterogeneous populations with distinct RMAE patterns. However, clonal sweeps can occur at the allelic level when physiological or pathological pressures selectively expand cells expressing particular allelic variants. For example, antigen exposure drives preferential expansion of lymphocyte clones expressing specific T- or B-cell receptor alleles^9^, while selective pressures imposed by therapy or immune surveillance can result in the dominance of tumor subclones carrying advantageous allelic variants^10,11^. X-chromosome abnormalities in particular have been implicated in the pathogenesis of basal-like breast cancers^12^. Loss of X-inactivation is a recurrent abnormality in breast cancer^13^, whereas studies in fibroblast and lymphoblast transcriptomes suggest that X-inactivation escape reflects a more complex biology observable only at single-cell resolution^14^. Moreover, potentially functional non-coding mutations in regulatory elements have been associated with significant differential allele-specific expression (ASE) between tumor and matched normal tissue, highlighting the need for allelic expression profiling for uncovering gene dysregulation in cancer^15^.

Breast cancer is a heterogeneous disease with molecular classification commonly guided by immunohistochemical (IHC) markers (ER, PR, and HER2) and transcriptomic signatures such as PAM50^16^, which stratify tumors into Basal-like, HER2-enriched, Luminal A, Luminal B, and Normal-like sub-types. While these frameworks provide prognostic and therapeutic value, several challenges remain for expression-based subtyping. Tumor heterogeneity and intermediate receptor expression (e.g., ER-low or HER2-low)^17,18^ can yield ambiguous assignments, and IHC- and transcriptome-based classifications are not always concordant. Moreover, current expression classifiers, derived largely from bulk tissue, may obscure intratumoral diversity and transitional states, limiting their resolution in capturing clinically relevant subgroups^19–21^. While the analysis of gene expression from large cohorts remains essential, we hypothesize that analyzing allelic gene expression provides molecular insights that uncover recurrent patterns otherwise missed in bulk sequencing analysis. Several analytical methods exist for calculating relative allelic gene expression in normal tissue^22,23^ from bulk and single-cell expression data; however, accurate quantification and inference of allelic expression in tumor tissue are hampered by varying tumor cell content in specimens as well as limited sensitivity for detecting somatic, focal copy-number changes that together confound measuring relative maternal and paternal allele expression in tumor cells while accounting for unaltered allele frequencies in normal cells in the specimen^24–28^. To address these challenges, we developed a novel quantitative method for accurately measuring imbalanced allelic expression in individual tumor samples using both genomic and transcriptomic data. We applied this framework to map the landscape of allelic imbalanced expression in breast invasive carcinoma and discovered a distinct subset of tumors, which did not align with previously defined gene expression-based molecular classifications, characterized by monoallelic expression of specific functionally relevant genes (Figure 1). Strikingly, this tumor subgroup, diagnosed primarily in patients of Black/African American ancestry and enriched in lobular tumors, was associated with significantly poorer overall survival relative to luminal tumors and showed comparably poor outcome to aggressive basal tumors. These results highlight the potential diagnostic and prognostic importance of monoallelic expression in this under-recognized tumor group and suggest new avenues for developing ancestry-informed biomarkers and therapeutic strategies.

**Figure 1.**
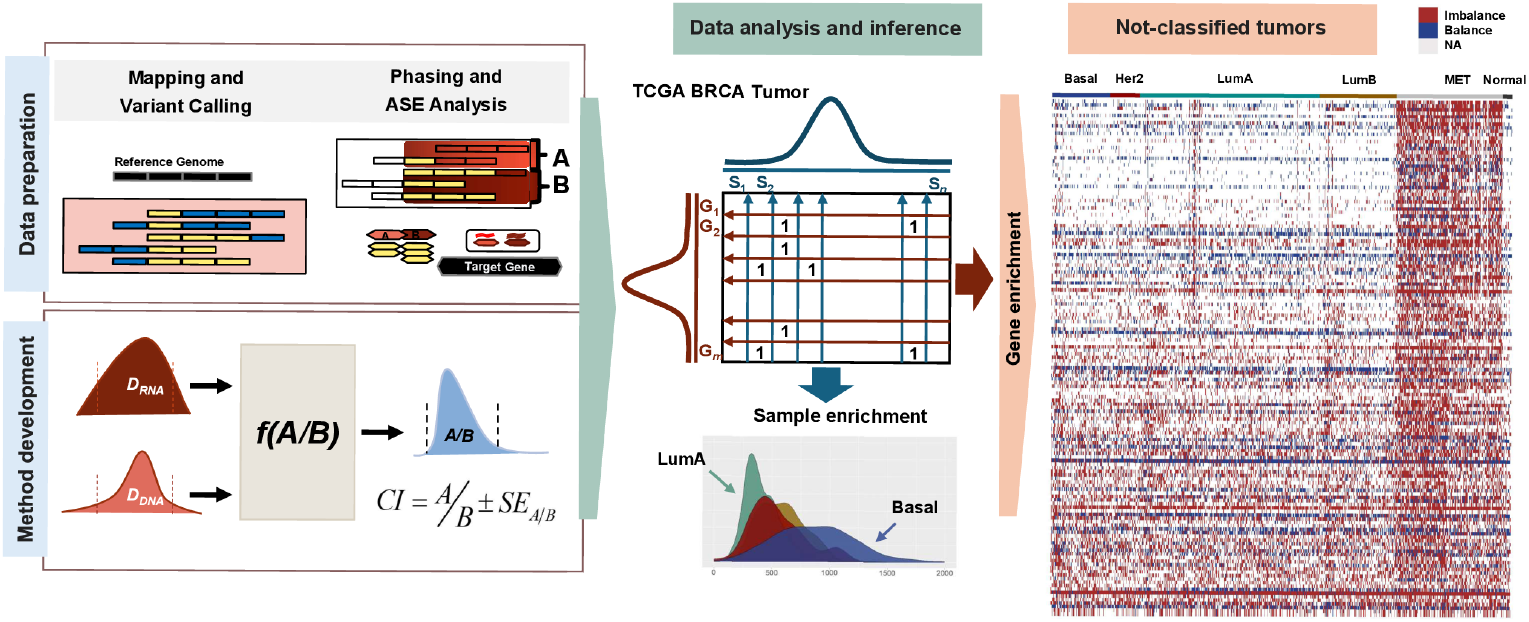
Schematic overview of the study design. The workflow includes data preparation, development of method for allelic imbalance detection, analysis of TCGA breast cancer tumors, and enrichment testing of allelic imbalance, highlighting distinct patterns in the unclassified tumors. Our workflow begins with data preparation, where we integrate tumor RNA-seq and DNA-seq data with matched germline variant calls to phase heterozygous SNPs and define informative loci for allelic expression analysis. For each gene, we derive allele-specific read counts from both DNA and RNA data. We then apply an interval-based allelic imbalance detection framework that accounts for expected variation in allelic depth and models the propagation delay inherent to read coverage along transcripts. This method improves discrimination between balanced and imbalanced expression states at the gene level. Then, we apply the method to TCGA BRCA tumors to assess, for every gene in every sample, whether its allelic expression is balanced or imbalanced. We then perform enrichment analyses at both the sample level and the gene level. Sample-level enrichment identifies tumors with disproportionately high numbers of monoallelically expressed genes, revealing subtype-specific patterns. Gene-level enrichment highlights sets of genes consistently exhibiting allelic imbalance across groups of tumors. Notably, these enrichment patterns delineate a subset of tumors that remain unclassified by PAM50, suggesting that allelic imbalance signatures capture biologically meaningful structure absent from conventional subtype assignments. Hereafter, we call this group monoallelic expression-enriched tumors (MET).

## RESULTS

### Interval-Based Allelic Imbalanced Detection Method

Accurate quantification of monoallelic versus biallelic gene expression in cancer cells requires a statistical comparison of relative allelic abundance in the transcriptomic and genomic sequencing data. To this end, we developed an Interval-Based Allelic Imbalanced Detection (IB-Aid) method that 1) corrects for tumor cell content (purity) and copy-number variations by normalizing allele frequencies measured in RNA using those in DNA and 2) accounts for sequencing biases by propagating variations in sequencing read counts (depth). IB-Aid calculates the confidence intervals (CI) for DNA-corrected, relative allelic expression and defines imbalance expression when the CIs reject a balanced expression hypothesis (i.e., the intervals around relative allelic expression do not include the ratio of one). IB-Aid is also able to evaluate relative allelic expression in non-tumor tissue samples where it is reasonable to assume a mean heterozygous germline variant allele frequency of 50% in DNA to which measured allele frequencies in RNA can be compared^29^.

We evaluated the impact of specimen tumor content, sequencing depth, and copy-number variations on the power to detect imbalanced expression by simulating 100 loci with different levels of tumor copynumber (2–10) in DNA with varying tumor purity (0.0–1.0) in samples. We considered germline and somatic mutant alleles independently and incorporated sequencing noise using binomial sampling^30^. We simulated imbalanced expression of either the mutant or the wild-type allele, and calculated specificity by simulating balanced expression of both alleles. Together these simulations allowed computing precision (or positive predictive value) as well as overall accuracy. As expected, the lower the tumor content, the higher the read counts needed to distinguish tumor cell-specific imbalanced expression from balanced expression of the same alleles in contaminating normal cells in the specimen. When samples with tumor contents *>*33% and sequencing read counts *>*50x were considered, overall sensitivity for correctly detecting imbalance expression was 0.76 with a precision of 0.97 while overall specificity for correctly detecting balanced expression was 92%. Overall accuracy and F-1 score were 0.80 and 0.85, respectively. There were no differences in overall performance when only either somatic or germline mutants were used except when purity was *≤*33% (Supplementary Figure 1).

### Tumor-Specific Patterns of Allelic Expression in Breast Cancer

Tumors that are associated with high levels of chromosomal instability also exhibit the highest degree of allelic imbalanced gene expression^31,32^. In breast invasive carcinomas (BRCA), particularly in aggressive subtypes like triple-negative and basal-like tumors, chromosomal instability is a hallmark, and it contributes to tumor evolution by driving copy number alterations and aneuploidy. These genomic changes often lead to allelic expression imbalances, such as loss of heterozygosity or monoallelic expression, affecting key tumor suppressor genes like *BRCA1* and *TP53*. These alterations can disrupt gene dosage and regulatory elements, leading to unequal expression of maternal and paternal alleles and promoting cancer progression and therapy resistance^33,34^.

To investigate tumor-specific patterns of allelic expression in breast invasive carcinoma, we used IB-Aid and analyzed DNA and RNA sequencing data from 934 tumor from patients with BRCA in The Cancer Genome Atlas (TCGA) (Supplementary Figure 2). The whole-exome DNA sequencing is limited in its scope to coding polymorphisms which may not be present in heterozygous genotypes across the patients in all genes. Moreover, DNA and RNA sequencing may lack sufficient read counts per allele per case, impeding the power to differentiate maternal and paternal allele frequencies. Therefore, we first evaluated the power for measuring allelic expression, and across the TCGA breast tumor samples, detected an average of 1.8 (range: 1–145) and 1.3 (range: 1–158) single nucleotide polymorphisms (SNPs) per gene per case in DNA and RNA sequencing data, respectively (Supplementary Figure 3). Moreover, the median total allele counts for each gene across all tumor samples was 147 (range 34–6057) for whole-exome data, which was significantly higher than the median total counts of 96 (range 20–50607) in RNA-seq data reflecting the uniformity of DNA sequencing depths in contrast to expected substantial variation in RNA read counts (Supplementary Figure 3). To attain consistency across the analyses, we used the genes that harbored ≥1 phased heterozygous polymorphic alleles and had sufficiently powered read counts per allele as a marker for robustly quantifying allelic expression. Therefore, we classified allelic expression as balanced (biallelic expression) or imbalanced (monoallelic expression) by computing allelic expression ratio and associated confidence intervals for 18,773 protein-coding genes, which identified 6,580 genes sufficiently powered to evaluate allelic expression in at least 10% of their samples (Supplementary Table 1). Among these genes, 88% showed imbalanced expression in less than 40% (Figure 2-A), while 7% showed recurrent imbalanced expression in more than 60% of their respective informative samples (Figure 2-B).

**Figure 2.**
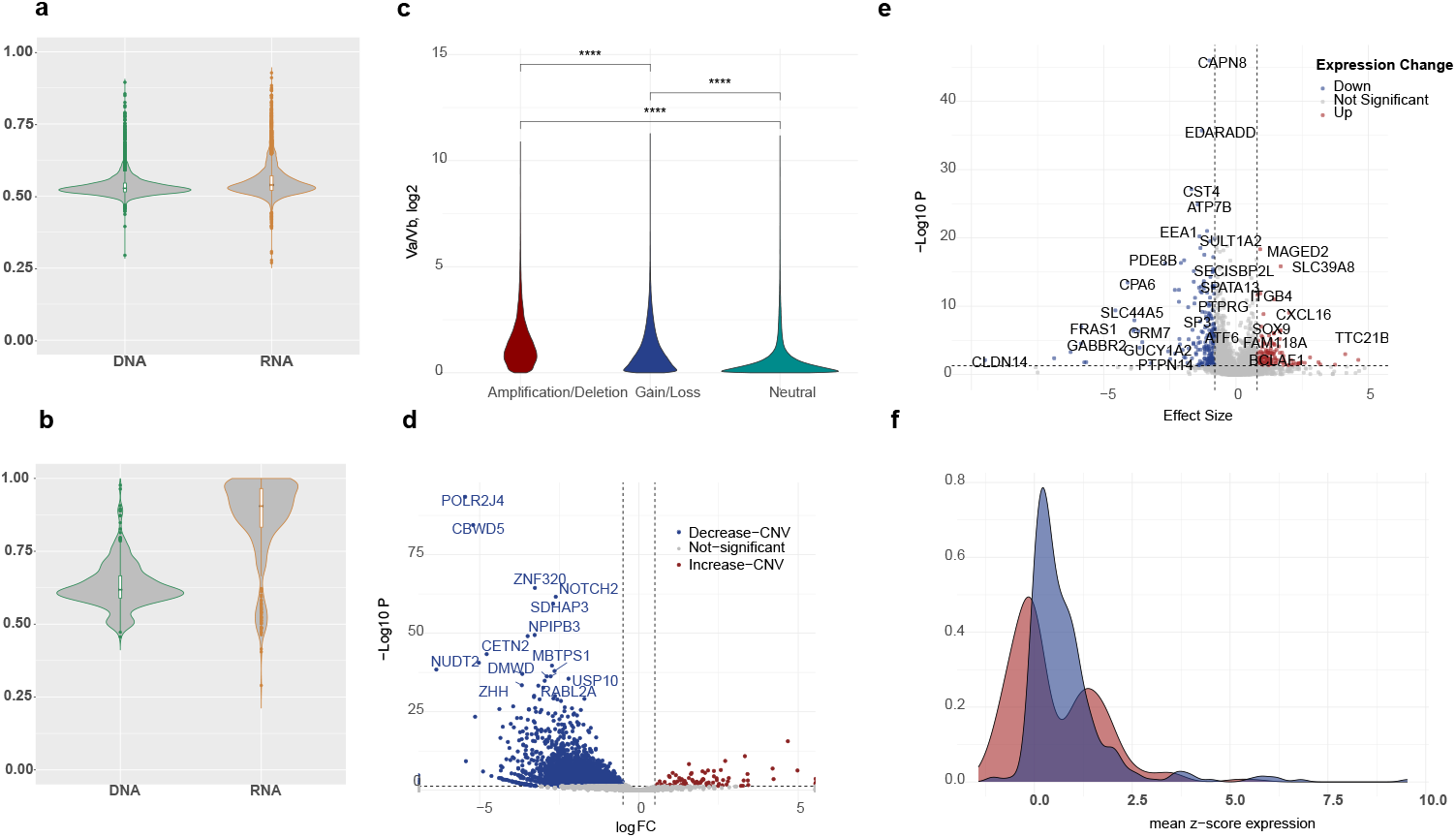
Allelic imbalance prevalence, copy-number contribution, and transcriptional consequences in breast tumors. A) DNA and RNA allele frequencies for genes with balanced allelic expression for 88% of 6,580 sufficiently powered genes. B) RNA and DNA allele frequencies for a subset of 7% of genes showed recurrent imbalance in more than 60% of their respective samples. C) Relative haplotype allele frequencies from DNA sequencing correlated with genome-wide copy-number gains and losses, demonstrating that allelic depth reflects underlying copy-number architecture. D) Correlation of allelic imbalance with copy-number variation, revealing that imbalance in approximately 50% of genes can be attributed to copy-number changes. E) Volcano plot of differential expression analysis between tumors with balanced and imbalanced allelic expression, indicating significant transcriptional changes in only 5.6% of genes. F) Distribution of mean z-scores for tumors with balanced versus imbalanced expression, showing two bimodal high and low expression for imbalanced samples.

#### Copy-number alterations underlie some but not all allele specific expression changes

To determine the impact of copy-number variations on allelic expression in breast tumors, we utilized the IB-Aid framework’s DNA-adjusted allelic transcription counts and compared allelic RNA expressions with and without correction for genomic copy-number. We first evaluated relative haplotype allele frequencies measured in DNA against the tumors’ genome-wide copy-number profiles. Relative total depth of DNA sequencing as well as observed relative frequencies of each two alleles correlated with presence of copy-number amplifications and deletions, including deeper copy-number gains and losses (Figure 2-C, Supplementary Figure 4); these results indicated that relative haplotype allele frequencies provide a quantitative measure that reflects underlying copy-number architecture and can be leveraged to refine detection of allele-specific copy-number imbalance across the genome. We further used these results to validate and normalize for copy-number variation in our method.

When we correlated imbalance expression with the presence of copy-number variations, we found that imbalanced expression in ≈ 50% of genes can unambiguously be attributed to copy-number changes in cancer cells (Figure 2-D, Supplementary Table 2). Basal tumors showed the most recurrent copynumber-associated monoallelic expression affecting 16% of evaluated genes (1,053 out of 6,580) followed by 10% of the genes in Luminal B tumors, 10% of the genes in Luminal A tumors, and only 0.4% in Her2-positive tumors. These findings indicate that the remaining fraction of imbalanced expression likely arises from transcriptional or epigenetic regulatory mechanisms independent of DNA copy-number alterations.

#### Allelic Imbalance can occur without changes in overall gene expression

We asked whether transcriptional profiles of genes with balanced versus imbalanced allelic expression were different. To address this, we calculated the differential mean gene expression per gene between the two tumor sets with balanced and imbalanced expression (Supplementary Table 3). We observed significant change in expression of only 5.6% genes (Figure 2-E.F). Transcriptional downregulation occurred in 224 of 6,580 of the genes (3.4%), including transcription factor genes such as *BRCA1, TP63*, and *SP1*; and groups genes belonging to the *TMEM, KRT* and *PTP* families. In contrast, 2.2% of imbalanced genes (143 of 6,580) showed increased expression including *MAGED2* and *CXCL16*^35–38^, possibly reflecting loss of imprinting or immune activity regulation. Gene-set enrichment analysis of these genes particularly showed involvement in apoptosis regulation (including *ITGB4, MYC, BCL2A1, BID*, regulation of apoptosis by parathyroid hormone-related protein, *P* = 0.007), and survival signaling (including *PFKL, BCL2A1, SERPINE1, BID*, Photodynamic Therapy Induced HIF 1 Survival Signaling, *P* = 0.02). The relationship between allelic expression and total expression is complex, as overall gene expression is typically assessed on normalized RNA-seq counts, whereas allelic imbalance captures deviations between the two alleles; this distinction raises the important question of when tumor biology depends on overall expression levels, associated primarily with underlying copy-number changes, and when it is driven by allelic gene imbalance, linked to regulatory mechanisms independent of DNA dosage.

### Prevalence of Monoallelic Expression and Genes Exhibiting Recurrent Monoallelic Expression across Breast Tumors Revealed by Sample-Specific and Gene-Specific Imbalance Enrichment

#### Prevalence of monoallelic expression across breast tumors

To investigate the distribution of monoallelic expression across breast tumors, we first evaluated monoallelic expression in individual tumor samples and found that those with the highest rates of imbalanced expression (i.e., top 5%) were classified as Basal tumors (*P* = 3.05e-05), whereas the lowest rates (bottom 5%) were of Luminal A subtype (*P* = 8.2e-10). These results suggest that enrichment of monoallelic preference at the sample level is subtype-associated and potentially reflective of changes linked to the extensive chromosomal instability in these subtypes and may serve as a reliable distinguishing molecular marker by itself (Figure 1).

#### Genes with recurrent monoallelic expression

To determine the genes most frequently affected by monoallelic expression across breast tumors, we evaluated recurrent imbalanced allelic expression across the entire cohort as well as within specific breast cancer molecular subtypes. This approach allowed us to identify both pan-breast-cancer events and subtype-restricted regulatory alterations that may contribute to tumor heterogeneity and clinical behavior. We identified a set of 241 genes that exhibited consistent monoallelic enrichment across the entire cohort, independent of tumor subtype (Supplementary Table 4). Notably, 3 out of 241 of these genes fall into well-characterized imprinted genes (*SNRPN, NLRP2, SNURF, ERAP2*-predicted), where expression is normally restricted to one parental allele due to epigenetic silencing; 13 are X-chromosome inactivation (XCI)-associated genes, where one X chromosome is transcriptionally silenced as part of dosage compensation (Supplementary Figure 5). Additionally, we observed monoallelic enrichment in eight autosomal genes (90% imbalanced in 90% samples of the cohort) not classically associated with imprinting or XCI, suggesting the involvement of potential tumor-specific regulatory mechanisms such as somatic epigenetic modifications, allele-specific chromatin accessibility, or noncoding variation affecting cis-regulatory elements (*BCLAF1, MAP2K3, PRIM2, PTPN14, NOMO1, AK2, AP3S1, TMEM128*) (Supplementary Figure 6). Finally, to identify subtype-specific patterns, we calculated the enrichment of monoallelic expression within each subtype (Supplementary Table 4). We stratified tumor samples by subtype using TCGA’s molecular PAM50 classification. We found the genes that exhibited significantly higher rates of monoallelic expression in one subtype compared to the rest of the cohort were enriched in the Basal and Luminal A/B tumors (Supplementary Figure 7-8), reflecting underlying biological differences in allelic control and possibly serving as candidate markers for further subtype-specific functional investigation.

### Characterizing Recurrent Monoallelic Genes in Unclassified Tumors

We used unsupervised analyses of samples across the whole cohort as well as supervised analyses of samples within each subtype and identified a group of tumors which showed exclusive enrichment in a subset of genes with recurrent monoallelic expression (Figure 3-A). These tumors were enriched in the group of samples which are not classified into conventional PAM50 subtypes. We found 145 genes (out of 6,580; 2%) that show recurrent monoallelic expression in these molecularly unclassified tumors marked by SNPs which were enriched in patients with these tumors relative to the rest of the cohort. Among the unclassified tumors, 46% were infiltrating ductal carcinomas (n = 100, enriched in Black/African American - hypergeometric test *P <* 0.005) and 42% were infiltrating lobular carcinomas (n = 91, enriched in White - hypergeometric test *P <* 0.005). In addition, 21% (46 of 216) of these tumors harbored somatic mutations in the *CDH1* gene, 93% of which were lobular carcinomas. Most strikingly, 47% (102 of 216) of patients with tumors enriched with monoallelic expression were Black/African American, in contrast to 17% (20 of 117) of patients with basal tumors (hypergeometric test *P* = 1.7e-35). Clinically, the unclassified, monoallelic expression-enriched tumors were associated with poor overall survival outcomes relative to Luminal A tumors (log-rank *P* = 0.09) while showing comparably poor outcomes to the aggressive Basal tumors (log-rank *P* = 0.05), Figure 4-A, suggesting a potential link between allelic imbalance and disease severity and outcome.

**Figure 3.**
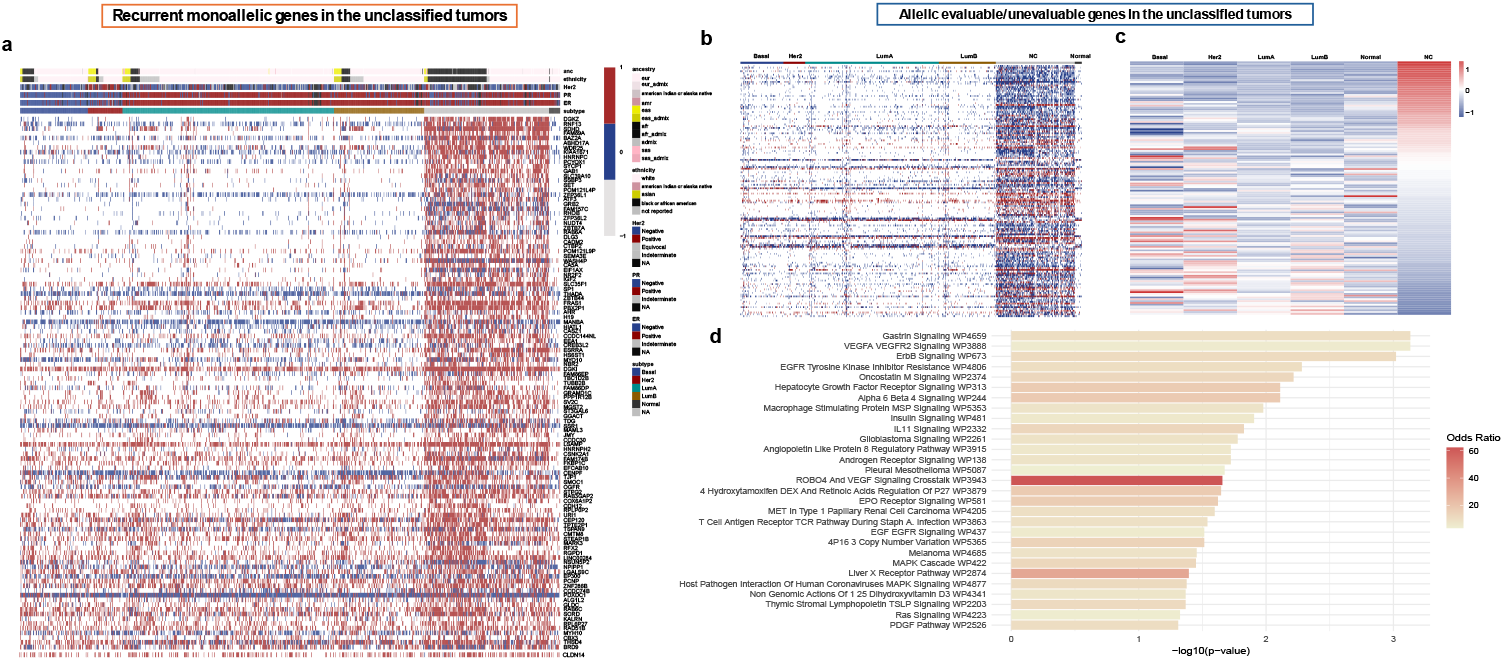
Allelic expression features distinguishing the MET breast cancer subgroup. A) Allelic expression patterns of a subset of genes showing monoallelic enrichment in the MET samples. B) Allelic expression patterns of genes specific to the MET samples with sufficient/insufficient read depth or informative SNPs to enable allelic expression evaluation. C) Expression profiles of the genes in panel B (shown in the same order), demonstrating that the MET samples cluster distinctly from other PAM50 molecular subtypes. D) Pathway enrichment analysis of genes in panel B, highlighting processes related to cellular signaling and metastasis, with key contributions from *MAPK3, GAB1, SHC2, SRC*, and *GRB2*.

**Figure 4.**
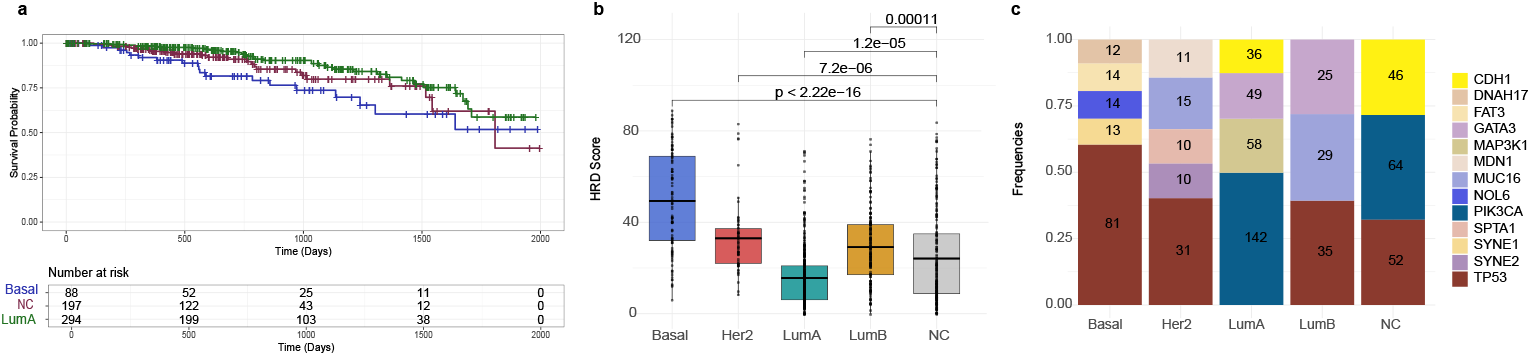
Molecular and clinical characterization of TCGA breast cancer subtype groups. A) Differences in progression-free survival between subtype groups. B) Median homologous recombination deficiency (HRD) scores across subtypes. C) Frequency of the top somatic mutations in each subtype group.

In addition to the allelic imbalance enrichment, we observed that from 6,850 genes that could be evaluated for allelic expression across the cohort, 154 genes had markers that were specific to unclassified tumors while 9 genes consistently lacked informative markers relative to the rest of the cohort (Figure 3-B, Supplementary Table 5). On clustering all tumor samples based exclusively on the expression patterns of these 163 genes, the unclassified tumors group together distinctly from other PAM50 molecular subtypes (Figure 3-C). This observation suggests two possible explanations: (1) the germline mutational landscape in the unclassified group may be different, or (2) the expression levels of these genes may be mechanistically higher—resulting in greater power to detect allele-specific expression—or conversely lower in other tumors, leading to insufficient power for such detection. The latter possibility merited further investigation of the pathways associated with these genes to understand the functional context and potential altered regulatory mechanisms in the unclassified group. Among the enriched pathways, pathways linked with cellular processes and metastasis involving key genes such as *MAPK3, GAB1, SHC2, SRC*, and *GRB2*, were of particular interest due to their roles in oncogenic signaling and cell motility regulation (Figure 3-D).

### Molecular Characterization of Monoallelic Expression-Enriched Tumors (MET)

To further characterize the monoallelic expression-enriched subset of breast tumors, we first analyzed molecular features linked to breast cancer biology. We kept tumors which were classified as other subtypes in their original classification. In addition to the allelic signature, we sought to determine whether other molecular features could explain this category and further confirm our observations.

#### Mutational burden and somatic mutation frequency

To map the patterns of genomic stability and clonal somatic evolution in the MET tumors, we evaluated homologous recombination deficiency (HRD) scores and somatic mutation frequencies. The MET tumors exhibited intermediate HRD scores—lower than those observed in Basal-like tumors, but higher than in Luminal A, suggesting partial impairment of homologous recombination repair (Figure 4-B). In terms of somatic mutation patterns, the MET subset resembled Luminal A/B tumors, with a relatively high frequency of *PIK3CA, CDH1*, and *KMT2C* mutations, and overall moderate mutation burden (Figure 4-C). This combination of genomic features supports the notion that the MET tumors occupy a distinct mutational landscape, characterized by both partial DNA repair deficiency and Luminal-like mutational signatures, which is not captured by standard subtype frameworks.

#### Patterns of gene expression

To characterize the MET samples based on their gene expression profiles, we selected genes with at least 0.3 z-score deviation relative to other classes of tumors. While a z-score deviation of 0.3 corresponds to a small effect size (i.e., 62nd percentile), even such modest shifts can be biologically meaningful when they are consistent and statistically supported. We identified 2,157 genes up-regulated and 1,050 genes down-regulated in the MET samples (Figure 5-A), with enriched pathways including those related to Cellular Processes, Signal Transduction Pathways (all are involved in cytokine signaling, development, and immune regulation) and Protein synthesis machinery (KEGG Ribosome pathway, showing upregulated) (Figure 5-B).

**Figure 5.**
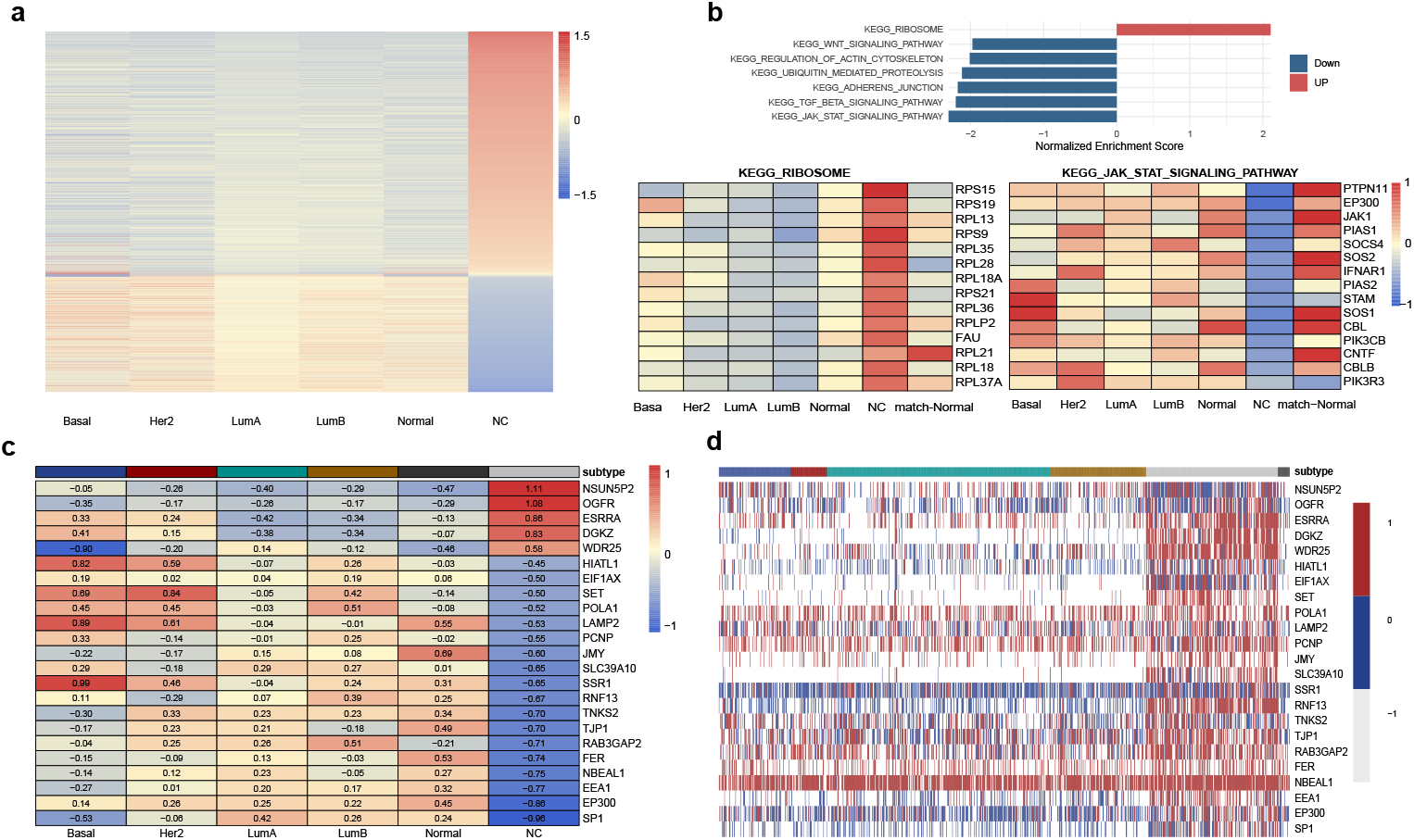
Transcriptomic and allelic expression features of the MET sample group. A) Differential expression profile of the MET tumors compared with other PAM50-classified tumors, highlighting genes showing significant up- or down-regulation. B) Enriched pathways in the unclassified group, including cellular processes, signal transduction pathways (notably cytokine signaling, developmental, and immune regulation), and protein synthesis machinery (e.g., upregulation of the KEGG ribosome pathway). C) Expression profiles of genes with differential expression between unclassified and classified tumors that also display allelic signatures. D) Allelic expression profiles of the genes in panel C, further illustrating allele-specific regulation distinguishing the unclassified group.

Among genes with differential expression profiles in the MET versus the PAM50-classified tumors, we found genes exhibiting both allelic and total differential gene expression, enriched in Adherens Junction pathway (*EP300, FER*, and *TJP1, P* = 0.003) (Figure 5-C,D). Within the MET samples’ gene signatures, *EP300* had low mean expression yet recurrently showed monoallelic expression. The central involvement of *EP300*, a known chromatin modifier, underscores the possibility that dysregulated allelic control in this group may be driven by altered epigenetic machinery. Moreover, the association of *EP300* across distinct sample clusters within this group suggests a unifying regulatory role, potentially contributing to shared transcriptional or epigenetic features that define this otherwise heterogeneous set of tumors.

Across the MET samples, we observed a distinct expression pattern marked by high *MGMT*, low *YY1*, and low *BRCA1* levels. *MGMT* overexpression suggests enhanced DNA repair capacity through direct reversal of alkylation damage, while reduced *YY1* —an *EP300* target—points to disrupted transcriptional regulation and chromatin remodeling. Notably, the low expression of *BRCA1*, a key gene involved in homologous recombination repair, may indicate BRCA pathway deficiency in this group. This constellation of expression features suggests a unique regulatory and DNA repair landscape in the unclassified group, potentially driven by epigenetic dysregulation and monoallelic expression events, with implications for therapeutic vulnerability to DNA-damaging agents and PARP inhibitors.

#### Cross validation of the MET group in the METABRIC dataset

To test whether the MET samples’ distinct gene expression profiles could be used to identify molecularly similar tumors in independent datasets, we trained a classification model using the TCGA data (based on a 5-fold cross-validation model with AUC = 0.993) and iteratively defined a set of differentially expressed marker genes that enabled discrimination between the MET samples from tumors belonging to other subtypes (Figure 6-A) in the METABRIC cohort^39^—a large, independent breast cancer dataset of 1,980 tumors with comprehensive clinical and molecular annotations. Through an iterative training approach, we identified 501 marker genes (Figure 6-B), Supplementary Table 6, which classified 224 tumors with MET-like expression profiles (Figure 6-C). These tumors were originally annotated across all PAM50 subtypes (38 among 329 Basal-like, 76 among 718 Luminal A, 27 among 488 Luminal B, 55 among 240 HER2-enriched, 25 among 199 Normal and 3 among 6 unclassified) and had survival outcome distributions similar to those seen in TCGA (Figure 6-D). These findings suggest that the MET samples define a novel transcriptionally distinct breast cancers that have been systematically overlooked by conventional classification approaches.

**Figure 6.**
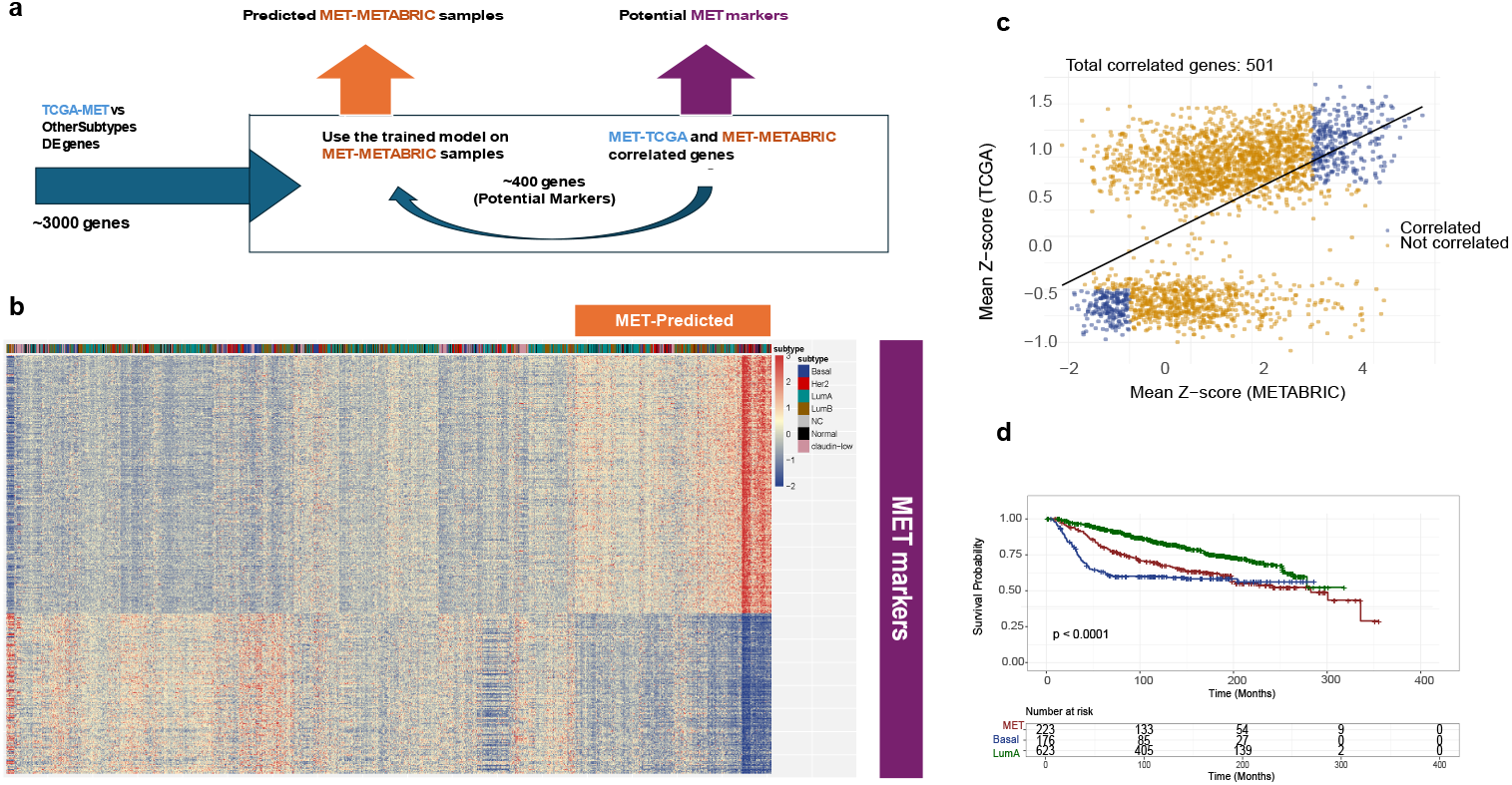
MET sample markers and subtype prediction across cohorts. A) Schematic framework for identifying marker genes of the MET sample group and predicting molecularly similar tumors in independent datasets. B) Genes with concordant expression patterns in both TCGA and METABRIC after the first and second training steps. C) Expression profile of samples identified as MET sample according to the classification model. D) Progression-free survival (PFS) differences between METABRIC patients of established subtypes and the predicted MET sample.

## DISCUSSION

Imbalanced allelic gene expression of both oncogenes and tumor suppressors have been implicated in development and progression of cancer. Cancer driver genes can exhibit imbalanced allelic expression due to mutations or other genetic alterations. For example, a copy of one of the alleles of an oncogene may be amplified or overexpressed, which can lead to an imbalance in the expression or activity of the oncogene. Similarly, tumor suppressor genes can undergo allelic imbalance due to mutations, deletion of one of the alleles, or other genetic alterations. Both alterations can promote the growth and proliferation of cancer cells. Thus, identifying and characterizing allelic imbalances in oncogenes and tumor suppressor genes can provide insights into the molecular mechanisms underlying cancer development and may help to inform the development of new cancer treatments. We showed that bulk sequencing eliminates enrichment of allelic expression across normal tissue and that tumor aberrations are associated with enrichment and co-exhibition of imbalanced allelic expression in genes involved in known and novel oncogenic processes. Copy-number variations may change both allelic expression and the overall expression of the genes. It is worth noting and reminding that the overall expression levels can be maintained through both monoallelic and biallelic expression, further highlighting the importance of additionally uncovering and understanding the role of monoallelic expression especially in the context of tumor development and progression.

The presence and the role of imbalanced variants in the marker genes of the MET samples group raise numerous questions that needs to be addressed in future studies; do variants linked to allelic expression contribute to tumor biology, is their imbalanced expression due to stochastic expression patterns, or can they be utilized as molecular markers associated with potential cancer drivers? Our results show that there are more allelically informative genes in the MET group of tumors—likely due to the presence of polymorphisms or informative SNPs—which may underlie their distinct allelic expression patterns. These findings further support a role for germline variation and allele-specific regulation in shaping subtype, and ancestry-linked gene expression in the MET samples. Some of these informative associations also show a clear link to ancestry, particularly among Black/African American patients. This is consistent with the ancestral enrichment observed within the MET subgroup, where these genes display allelic patterns shaped by population-specific SNP variation. The overlap between ancestry-associated polymorphisms and allelic expression profiles underscores the importance of considering genetic background when interpreting monoallelic expression, especially in transcriptionally unclassified tumors. Analyzing top phased polymorphisms within each gene to examine their associations across tumor subtypes and ancestry groups, stratified by balanced and imbalanced expression states, led us to SNPs which were consistently associated with the MET samples under both conditions, indicating their role as informative markers of allelic expression in this molecular subgroup. In contrast, we uncovered polymorphisms linked exclusively to either balanced or imbalanced expression, suggesting their potential functional or regulatory roles. Consistent with the enrichment of genetic backgrounds in the MET sample, several polymorphisms were ancestry-specific and present in the Black/African American patients, reinforcing the interplay between germline variation, allele-specific expression, and population background in shaping transcriptional phenotypes. For example, our analysis showed that in the SDHD gene, the absence of two variants in both DNA and RNA transcripts in the tumor resulted in the formation of an absent haplotype, whereas the germline variants associated with the MET samples were present in the matched normal tissue (Supplementary Table 7). The potential role of these variants in driving allelic imbalance —and whether such imbalance may serve as a molecular marker for this group—remains an open question under active investigation.

The concurrent low expression of *BRCA1* and *EP300* in the MET samples suggests a potential functional link between chromatin regulation and homologous recombination repair deficiency; the downregulation also consistently has been observed in the *CEP170* gene, also known as centrosomal protein 170, which is involved in several cellular processes, including microtubule organization and double-strand break repair, and has been linked to cancer progression^40^. Specifically, research suggests that mutations or low levels of *CEP170* in various cancer types, including breast cancer, can impact cell polarity, motility, and response to DNA damage. *EP300* is a histone acetyltransferase that facilitates chromatin accessibility and serves as a transcriptional co-activator for numerous genes, including those involved in DNA damage response^41,42^. Its reduced activity may impair the transcriptional activation of *BRCA1*, directly or indirectly, thereby contributing to BRCA pathway suppression. This regulatory connection is supported by prior studies showing that *EP300* can influence *BRCA1* promoter activity through histone acetylation and chromatin remodeling. Thus, *EP300* deficiency may play a mechanistic role in the observed *BRCA1* downregulation, reinforcing the possibility of epigenetically driven BRCA deficiency in this subgroup and highlighting potential sensitivity to DNA repair-targeted therapies^43,44^.

Among the genes linking allelic expression, transcriptional count, and clinical outcome, *MAGED2* and *CAPN8* emerged as key markers. *MAGED2* showed higher expression in balanced samples, indicative of biallelic expression and potential loss of X-inactivation, while *CASP8* displayed lower expression in imbalanced samples. Notably, balanced expression of *MAGED2* and imbalanced expression of *CAPN8* were both associated with shorter survival, suggesting a link between allelic status and clinical progression. Functionally, *MAGED2* has been shown to act as a negatively regulator of *TP53* activity, and its loss is associated with metastatic phenotypes in breast cancer^36,45^. These findings support the role of allele-specific regulation in modulating key tumor suppressive and prometastatic pathways. In our analysis, *CAPN8* emerged as a gene showing allelic imbalance associated with reduced expression in a subset of tumors. While primarily characterized in gastric and thyroid cancer, its role remains underexplored for breast cancer and warrants further investigation^46,46^. These findings underscore the prognostic relevance of allelic expression and support its potential as a stratifying biomarker, with allelic expression serving as a useful reference for evaluating clinical risk.

Short RNA sequencing reads constrain the power to phase SNPs and accurately quantify allelic counts, resulting in a reduced set of genes with phased counts available for both RNA and DNA analysis. Furthermore, while adjusting for DNA copy aims to account for somatic CNV, imbalanced RNA expression can still arise from extreme copy-number changes reflecting true imbalance beyond dosage. Despite these challenges, bulk-level analyses remain essential, as they capture genome-wide patterns across large cohorts and enable identification of recurrent allelic phenomena. Building on such discoveries, single-cell approaches can then be applied to specific tumor groups—such as those identified here—to disentangle cellular heterogeneity, refine mechanistic interpretation, and evaluate how allelic regulation contributes to tumor progression and therapeutic response.

## Methods

### Patient data

We retrieved The Cancer Genome Atlas (TCGA) raw RNA sequencing data for tumor and normal samples, as well as protected germline calls detected by DNA sequencing of normal samples^47^ for 934 patients with breast cancer from the GDC archives following an approved dbGaP project. We obtained TCGA clinical and molecular data from the Xena portal^48^. We obtained access to the METABRIC cohort gene expression and metadata for 1980 samples^39^, after formally requesting and receiving permission from the data custodians.

### RNA-seq and allelic expression analyses

We used STAR Ver 2.7.5a^49^ and GATK Ver 4.2.2.0^50^ best practices approach to align the RNA sequencing reads to the hg19 human reference genome, and used phASER^29^, a haplotype phasing method, to refine allele-specific expression by measuring the number of reads belonging to each allele (mapq and baseq were set to 0 for higher flexibility). We used the DEseq for normalization^51^ of raw expression counts and calculated the z-scores for down-stream expression level comparisons as previously described.

For the TCGA breast cancer cohort, we observe that on average, 24% of genes in each sample have phased SNPs and 30% of samples have phased SNPs for each gene. We also normalize the number of SNPs by the gene length and filter out genes that have the normalized values less than 2.5 standard deviation from the mean of the distribution. This process left 85% genes for our downstream analysis providing sufficient statistical power to arrive at reliable estimates for the allele counts. We used the same approach for the normal match samples with mRNA available data, and ended up with 85 normal samples.

### Interval-Based Allelic Imbalanced Detection (IB-Aid)

We developed IB-Aid to analyze both DNA and RNA data from the same specimens. IB-Aid corrects for purity and copy-number variations by normalizing AF in RNA using AF in DNA and computes allelic imbalance ratio and its confidence interval (CI) using uncertainty propagation to model technical sources of error (Eq. 1). IB-Aid is designed based on principles in Interval arithmetic, a technique used to put bounds on measurement errors in mathematical computation; therefore, instead of a single number, a range of possibilities are considered for a measurement. In our analysis the depth of RNA (*D*_RNA_) and the depth of DNA (*D*_DNA_) are experimental measurements, and we consider binomially distributed variant allele frequencies measured in RNA (*V*_RNA_) and in DNA (*V*_DNA_). The relative expression of allele A and allele B is then defined as Eq. 2. We propagate measurement errors for (*V*_RNA_) and (*V*_DNA_) to calculate confidence interval for the ratio of allelic imbalance^52^; Eq. 3-Eq. 5. The final confidence interval for allelic expression assess by replacing Eq. 2 and 5 in Eq. 1. When *V*_RNA_,*V*_DNA_=1 or *V*_RNA_,*V*_DNA_=0 or *V*_DNA_ = 0 or *V*_RNA_ = 1, we consider the ratio as infinity (a constant number 100), and *SE* =0. In Eq. 4, we calculate the Wilson score^53^ with 5% confidence interval with z=1.96. We define AI when the confidence interval does not coincide with balanced relative expression.

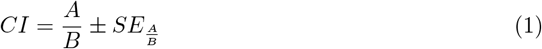

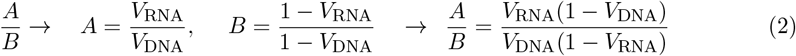

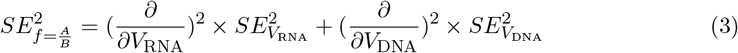

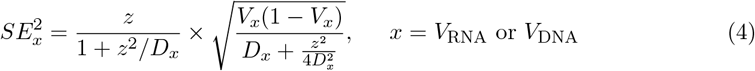

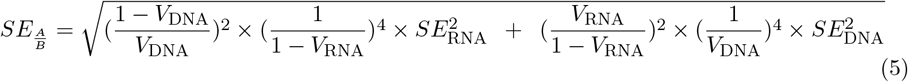

### Benchmarking IB-Aid

We evaluated the impact of specimen tumor content and the extent of copy-number variations on the power to correct for DNA impurities while assessing allelic imbalance using simulations. We used a consistent simulation model to describe germline and mutational status with the corresponding to the number of mutated copies for each mutation56. We performed analyses using simulations with somatic or germline mutations across purity (0–100%) and somatic copy-number (*>*2–10) values for one balanced dataset and three imbalanced datasets. We found that sensitivity for correctly detecting allelic imbalance was 0.76 with a precision of 0.97; specificity for correctly detecting the absence of allelic imbalance was 0.92 with accuracy and F-1 score of 0.80 and 0.85, respectively. There were no differences in overall performance when only either somatic or germline mutants were used except when purity was *>*33%.

### Application of IB-Aid to TCGA and GTEx

We calculated allelic imbalance of 18,773 protein-coding genes, for 934 TCGA tumor samples, 85 TCGA normal samples and 359 GTEx breast tissue samples, with the interval cut-off of 0.2 and 5 on the ratio of allele fraction. Therefore, when the standard error low-bar/high-bar pass 0.2/5 we tag the gene-sample loci in the matrix as imbalance. We applied a depth exclusion limit of *<*50 for either RNA or DNA considered to be as non-informative. The final n*×*m Allelic Imbalance Matrix for the dataset with n genes and m samples is a quad-value matrix; 0 balance, 1 imbalance, 1 not informative, 2 no-counts. Informative sample counts (having counts/informative AI calculation) which we will refer to as informative or sufficiently powered in the text, is the number of balanced samples plus the number of imbalanced samples for each gene.

### Statistical and survival analyses

The exact test was used to evaluate co-occurrence of mutations across the gene pairs. Statistically significant differences between groups were calculated for continuous or categorical variables using the exact test, chi-squared, or non-parametric rank-sum test as indicated. The Benjamini-Hochberg false discovery rate (FDR) correction was used for multiple hypotheses testing when indicated.

Clinical associations were determined using a multivariable Cox model regressing clinical values against different profiles. Statistical analysis and data visualizations were performed using following packages in R: surminer, survival (survival analyses), and ggplot2, pheatmap (plotting).

## DATA AND CODE AVAILABILITY

No original dataset was created as a part of this study. The IB-Aid method is implemented in R available at https://github.com/marabzadeh/IBAid.

## ACKNOWLEDGMENT

The authors would like to thank Drs. Chang Chan and Gregory Riedlinger, for careful reading of the manuscript and for providing constructive feedback.

## SUPPLEMENTARY INFORMATION

**Supplementary Table 1**. List of sufficiently powered genes to evaluate allelic expression in at least 10% of their samples.

**Supplementary Table 2**. Copy-number attributed genes and their subtype associations.

**Supplementary Table 3**. List of genes with differentially expressed profiles between their balance and imbalance samples.

**Supplementary Table 4**. List if recurrent monoallelic genes overall and subtype specific.

**Supplementary Table 5**. Genes sufficiently powered to be evaluated for allelic expression specific to the unclassified tumors.

**Supplementary Table 6**. Marker gene set for the unclassified group across cohorts.

**Supplementary Table 7**. Attributions of the top germline variants phased in the recurrent monoallelic genes of the unclassified tumors.

**Supplementary Table 8**. Specifications of X-chr genes in tumor and match-normal samples of TCGA.

**Supplementary Note**. Detailed description of additional information.

