## Supplemental Notes for "Monoallelic expression characterizes a distinct molecular and clinical group of breast tumors"

### Supplementary notes

***Interval-Based Allelic Imbalanced Detection Method Simulation.*** In samples with high tumor content >0.67, all performance measures were >0.85 when total read counts were >20x, reaching to >0.95 at counts above 50x. For tumor samples with purity in the mid-range of 0.33–0.67, sensitivity and accuracy reached to >0.85 when total read counts were >100x. In samples with very low tumor contents of ≤0.33, the accuracy for detecting imbalanced expressions reached to only 0.68 at depth >200x.

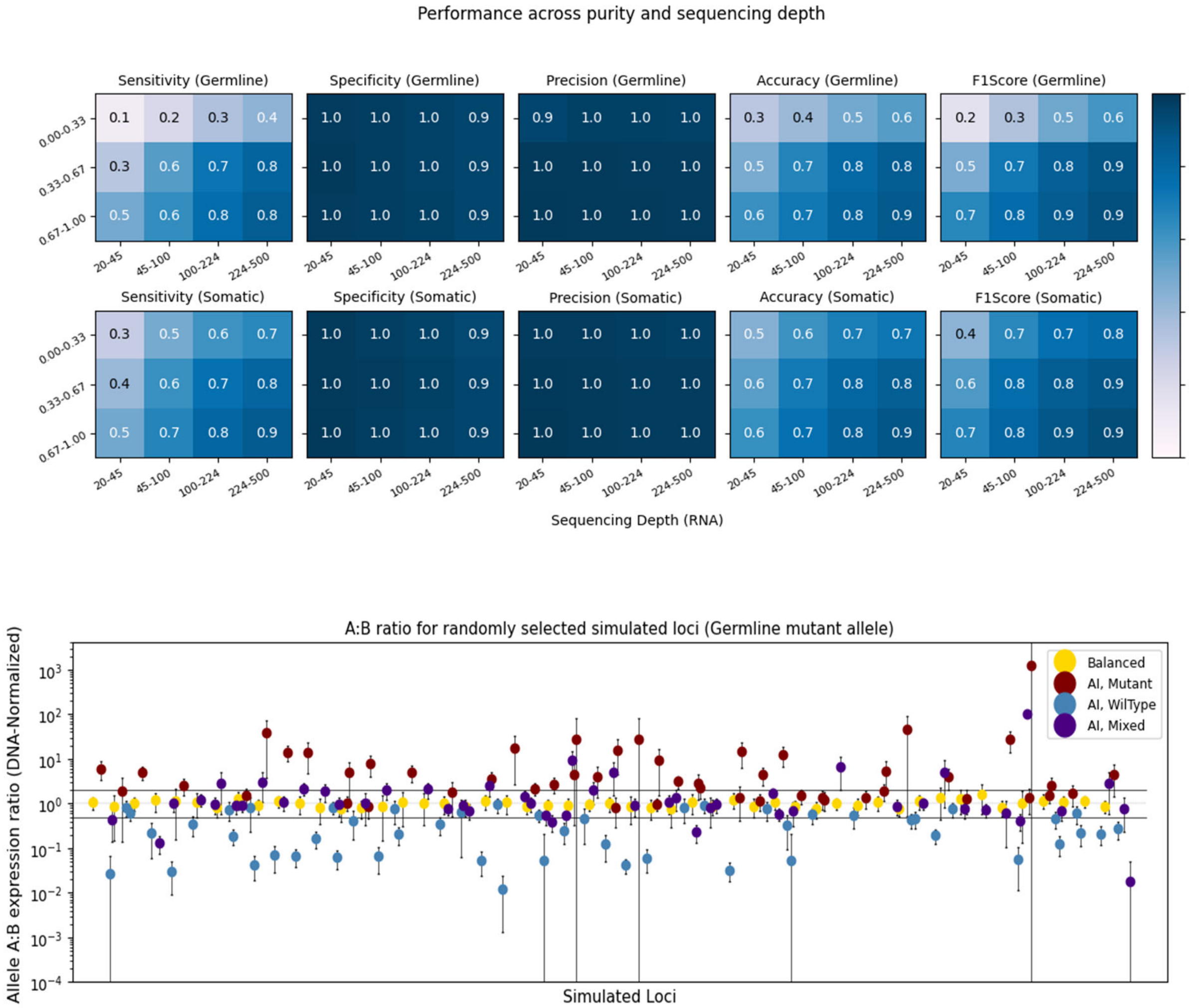


**Figure 1**. Allelic Imbalance detection method simulation results. A) Performance across purity and sequencing depth. B) A:B ration for randomly selected simulated loci.

##### *Framework to process the allele counts.*

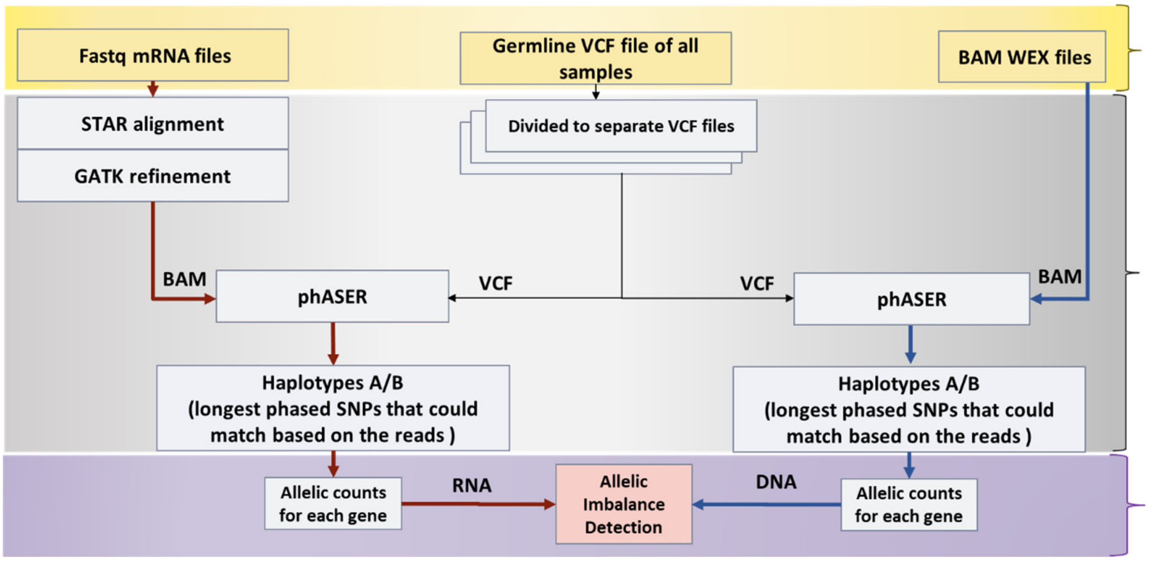


**Figure 2**. Framework that has been used for extracting allele counts for TCGA BRCA data.

##### *SNP and depth for TCGA tumor for sufficiently powered genes.*

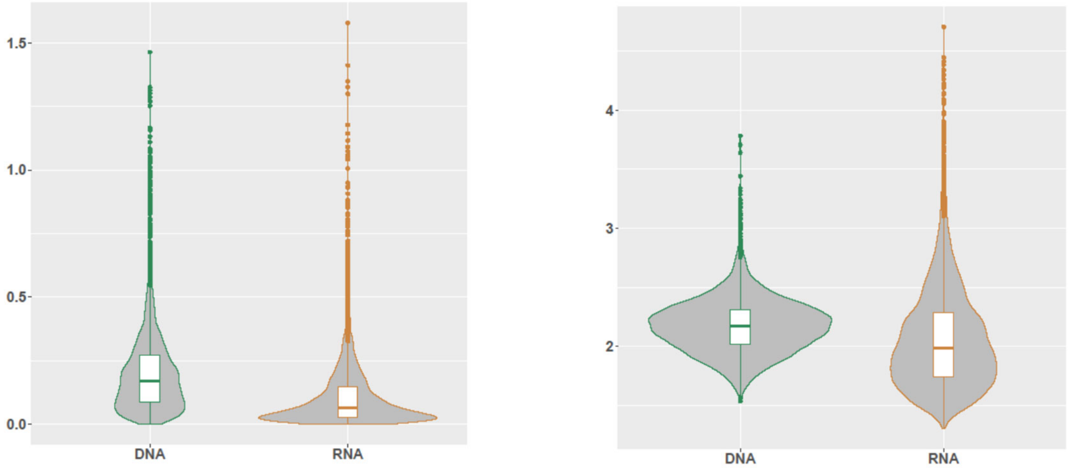


**Figure 3.** The distribution of mean SNP and depth for sufficiently powered genes for both DNA and RNA counts.

##### *Copy-number distribution of allele frequency in CNV profiles.*

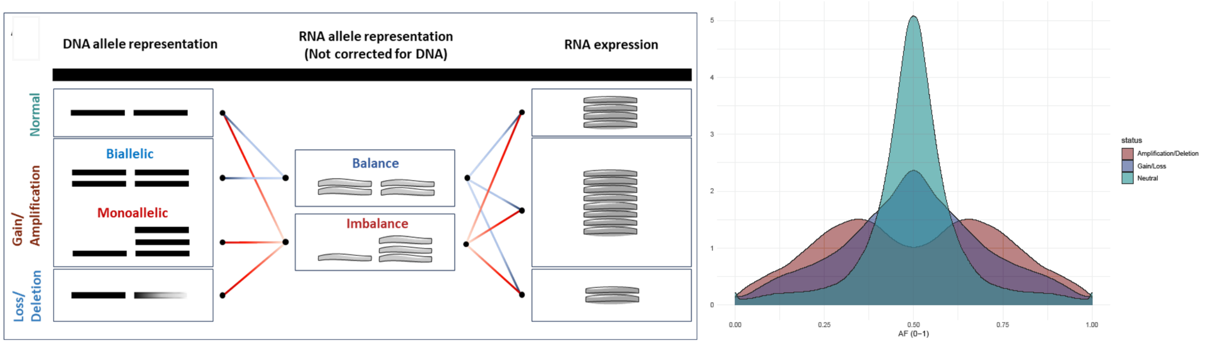


**Figure 4.** CNV and expression attributions. A) Schematic that showing how DNA copies, RNA allelic expression and overall expression can be related. All possible combinations should be considered while interpreting the results. B) Distribution of the allele frequency of DNA counts in different copy-number profiles calculated for TCGA data.

##### *Recurrent monoallelic genes.*

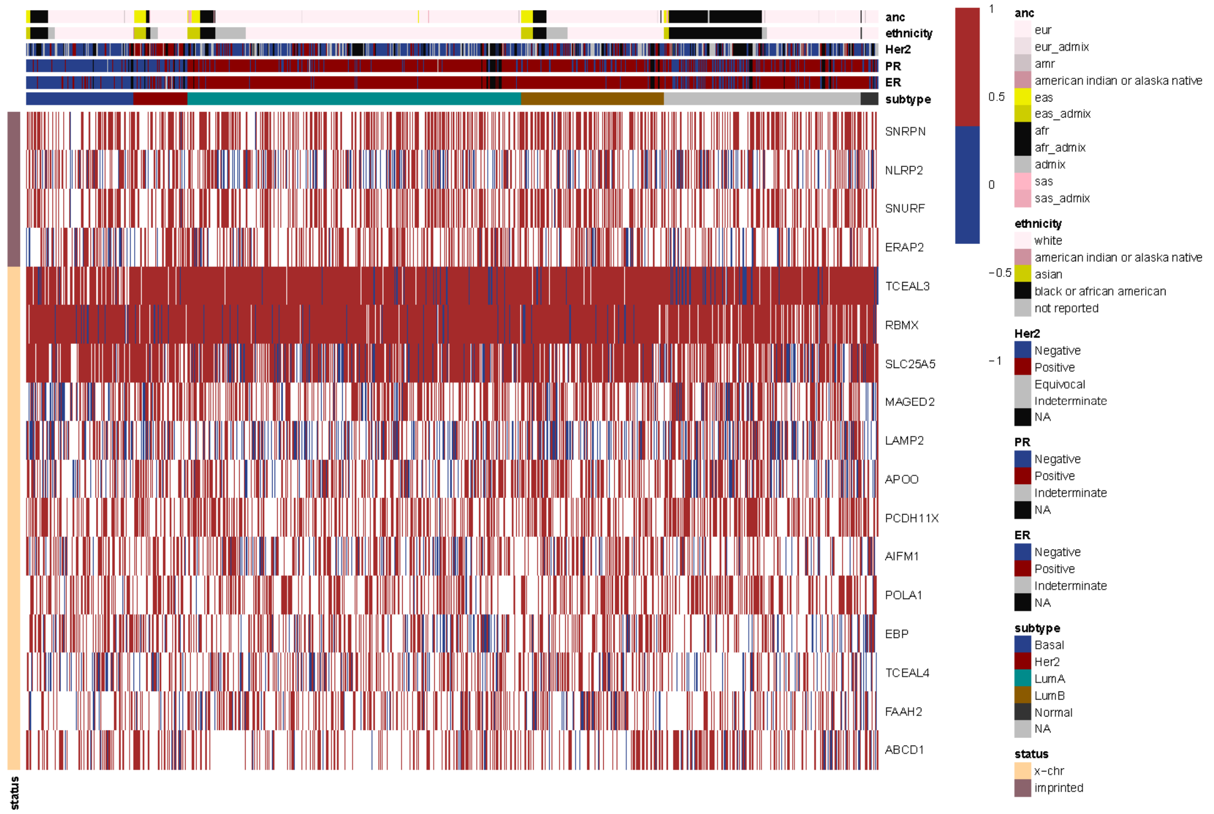


**Figure 5.** Allelic expression profile of x-chr and known imprinted genes.

Among the gene with the highest rate of imbalanced expression, we identified a subset—including BCALF1, MAP2K3 and AP3S1 —whose allelic expression is consistently restricted to a single allele across clonal tumor cells. These patterns suggest potential imprinting-like regulation, where gene expression is monoallelic due to inherent epigenetic programming or tumor-specific silencing mechanisms. Whether through parent-of-origin imprinting, epigenetic silencing, or allele-specific regulatory disruption, these genes appear to maintain stable monoallelic expression in the tumor context, raising the possibility that they contribute to clonal identity or tumor progression.

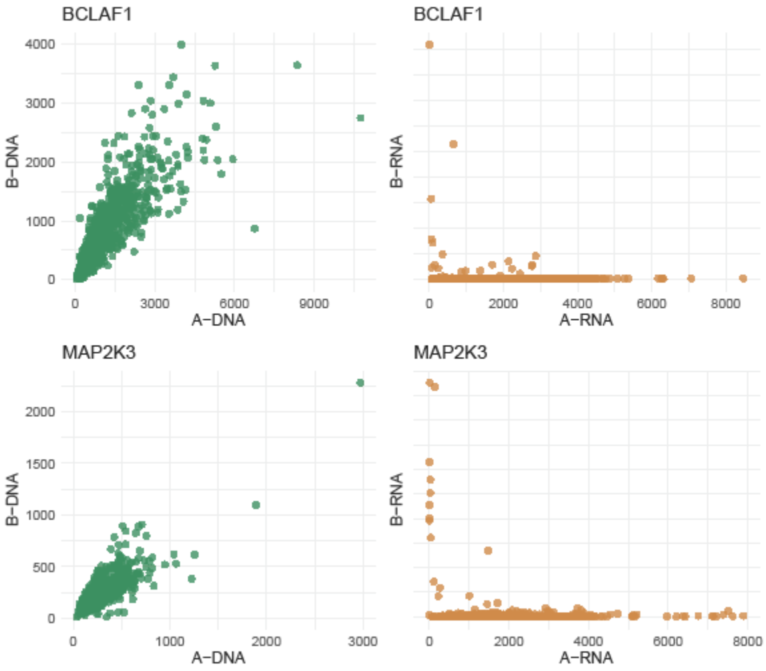


**Figure 6.** DNA And RNA allelic distribution of BCALF1 and MAP2K3.

***Subtype-specific genes pathway enrichment.*** Pathway enrichment analysis of subtype-specific monoallelic gene groups revealed distinct functional signatures reflective of each tumor subtype's biology. In the Basal subtype, genes exhibiting enriched monoallelic expression were significantly associated with *Keratin type II family members*, as well as pathways related to *cell cycle regulation* (adjusted $p$ = 0.006) and *DNA repair*, specifically the *Fanconi anemia pathway* (adjusted $p$ = 0.03). This suggests a potential link between allelic imbalance and critical processes involved in *proliferation* and *genomic stability* within basal-like tumors. In contrast, the LumA/B subtype showed moderate enrichment for pathways involved in *phospholipid biosynthesis* (adjusted $p$ = 0.09), *formation of the translation preinitiation complex* (adjusted $p$ = 0.09), and *pantothenate and CoA biosynthesis* (adjusted $p$ = 0.30), suggesting potential alterations in *membrane remodeling*, *translational control*, and *coenzyme metabolism* that may distinguish this subtype at a functional level. These distinct pathway enrichments suggest that monoallelic expression in each subtype may be shaped by—or contribute to—subtype-specific regulatory programs and may influence therapeutic responsiveness or tumor progression in a subtype-dependent manner.

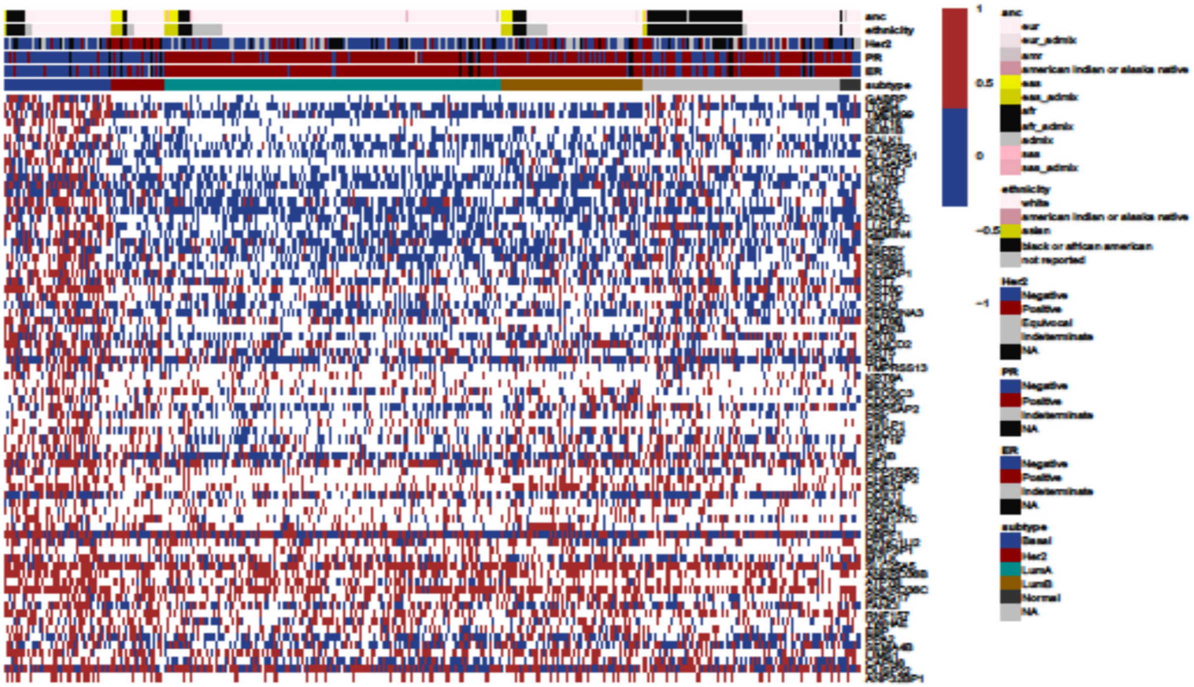


**Figure 7.** Allelic expression profile of recurrent monoallelic genes in Basal.

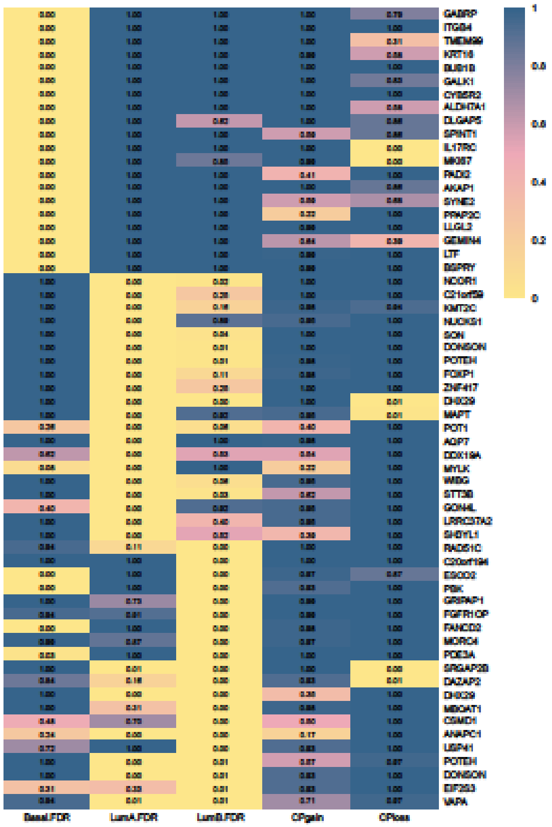


**Figure 8.** Top 10 recurrent monoallelic genes in each subtype, their exclusivity, and their copy number associations.

***Allele-specific expression (ASE) of pioneer transcription factors.*** ASE of TFs have been associated with disruption of regulatory networks and tumorigenesis. Among the **6,580 genes** tested for allelic expression, we identified 307 transcription factor genes, 8% of which (30 of 365) exhibited allelic imbalance in more than 50% of tumors. Therefore, we asked whether imbalanced expression of TF genes had an impact on expression of their target gene. To this end, we stratified the tumors into two groups with TF balanced and imbalanced expression of each factor and evaluated the expression levels of their known target genes by looking at the mean expression level between the two groups. We identified significant differences between expression of targets genes for 10 out of 30 recurrently imbalanced TFs, including *MYC*, *JUN*, *SP1*, and *ELF1*.

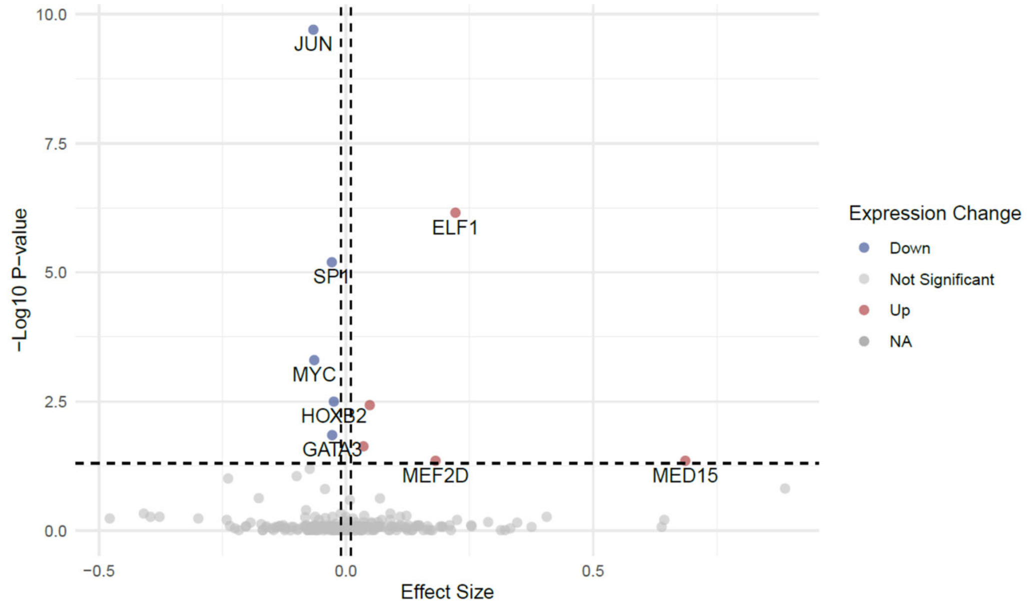


**Figure 9.** Volcano plot for differential expression profiles between balance and imbalanced samples of TF genes.

We classified the breast tumors based on the allelic expression profiles across these transcription factors and identified the factors naturally grouped together, forming patterns that separated specific subsets of tumors. There are two groups that are characterized by imbalanced expression of functionally related genes, including *NCOR1*, *KMT2X* and *ARID1A*, *BRCA1*, *CNOT7*, *NR3C1*.

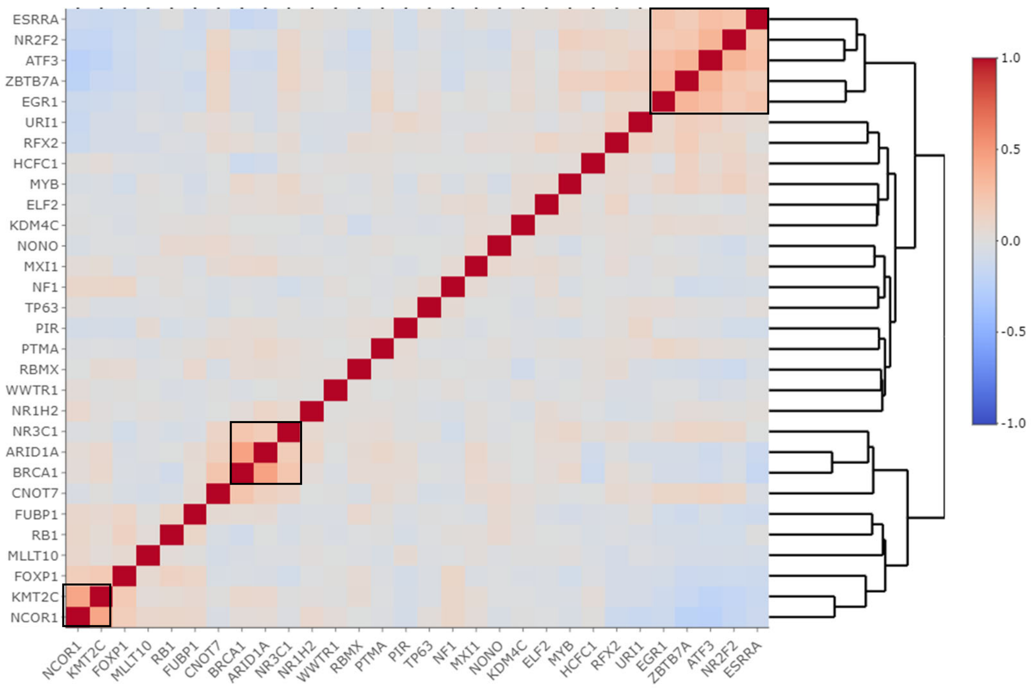


**Figure 10.** Correlation of imbalanced samples of TF genes. Higher correlation score indicates the more imbalanced samples of each two gene appear together.

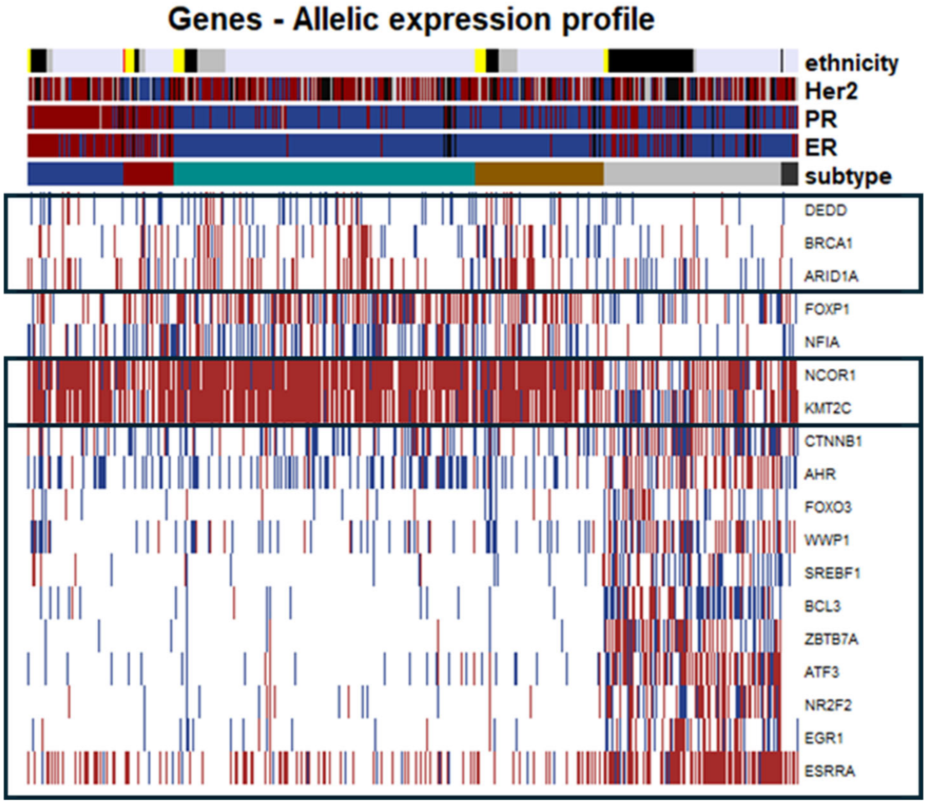


**Figure 11**. Allelic expression profile of correlated TF genes.

***Imbalanced expression associations with clinical outcome.*** Survival analysis showed that monoallelic versus biallelic expression of 113 genes (with more than 10 samples in each allelic profile group) was significantly associated with differences in clinical outcomes. Imbalanced expression was generally associated with poor overall survival (Give range in risk per time units) compared to balanced expression (in 110 cases out of 113 cases). Five genes, including LLGL2 (enriched in Basal), MLEC (enriched in LumA), TDG (enriched in not-classified), CAPN8, and ABCC8 showed worse survival outcomes in tumors with imbalanced, suggesting that allelic imbalance in these loci may contribute to more aggressive disease biology. In these genes, the average ratio between the number of balanced and imbalanced samples is 1.8. Conversely, imbalanced expression of a smaller subset of genes, including MAGED2, SET, and ITGB5 was associated with improved survival, indicating possible context-specific protective effects or immune-related mechanisms. Among these genes, MAGED2 showed monoallelic enrichment in Basal, and ITGB5 in the not-classified tumor samples, respectively. In SET and MAGED2, the average ratio between the number of balanced and imbalanced samples is 1.5, being dominated in imbalanced samples.

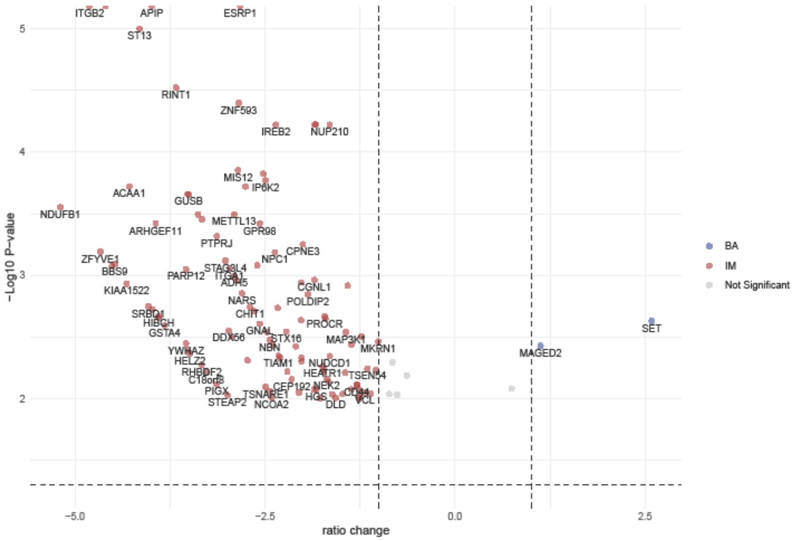

**Figure 12.** Volcano plot showing differential survival ratio and p-value between balance and imbalance samples of each gene.

***Comparison to normal tissue.*** We examined genes known as imprinted/monoallelic genes which exhibit balanced allelic expression in a subset of tumor samples in sharp contrast to the expected monoallelic exhibited in normal tissue (GTEx as reference). For instance, MEST and IGF2 are imprinted genes, which clearly lose imprinting in at least 15% of samples in TCGA-tumor. The loss of imprinting of MEST and IGF2 was not associated with any breast cancer subtype. Next, we examined whether there were genes which exhibited monoallelic expression in a significant fraction of samples both in the tumors as well as normals. These genes would reflect imprinting which has largely been preserved even in the tumors. With the threshold of 50% imbalance, for both GTEx and TCGA, we found RPL9 with strong evidence of monoallelic expression, has been reported to exhibit monoallelic expression.

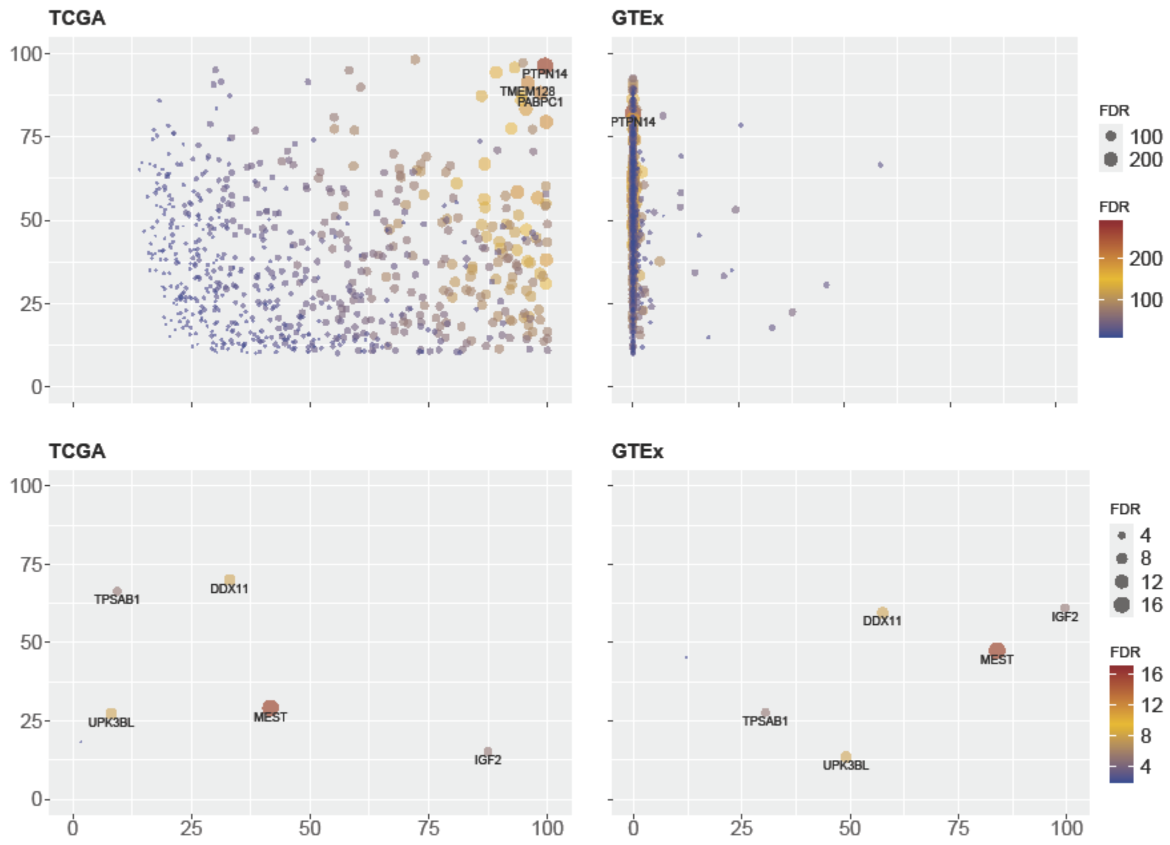


**Figure 13.** Genes showing highly differential allelic ratio in TCGA tumor and GTEx allele counts based on the balance/imbalance assessment with IB-Aid method. Y-axis indicates that in what percentage of samples each specific gene was sufficiently powered, and X-axis indicates that in what percentage of samples that specific gene showed imbalanced expression; looking at this information in both TCGA and GTEx (left and right). Top plots are genes with highly differentiated genes between TCGA and GTEx weighed towards being imbalanced in TCGA, and bottom plots are showing differentiated genes weighted towards being balanced in TCGA (loss of imprinting or X-chr inactivation).

When comparing the prevalence of monoallelic expression between tumors and matched normal samples, we observed some X chromosome genes displayed striking differences: while these genes showed a high percentage of allelic imbalance in tumors, the imbalance frequency was approximately halved in normal tissues **(Supplementary Table 8)**. This pattern may reflect dysregulation of X-chromosome inactivation or escape from dosage compensation in tumor cells, leading to preferential expression of one allele. Such alterations could contribute to sex-linked vulnerabilities in breast cancer and highlight a mechanism by which tumor cells exploit allelic imbalances.

***Gene-variant allelic and ancestry association.*** Among the exclusive genes in the unclassified group, part of genes showed strong association with allelic imbalance, while other were associated with both balanced and imbalanced states. This dual association suggests that these genes are allelically powered in this group—likely due to the presence of polymorphisms or informative SNPs—which may underlie their distinct expression patterns. These findings further support a role for germline variation and allele-specific regulation in shaping subtype- and ancestry-linked gene expression in the unclassified group. Some of these informative associations also show a clear link to ancestry (p-values), particularly among Black/African American patients. This is consistent with the ancestral enrichment observed within the unclassified subgroup, where these genes display allelic patterns shaped by population-specific SNP variation. The overlap between ancestry-associated polymorphisms and allelic expression profiles underscores the importance of considering genetic background when interpreting monoallelic expression, especially in transcriptionally unclassified tumors.

We further analyzed the top phased SNPs within each gene to examine their associations across PAM50 subtypes and ancestry groups, stratified by balanced and imbalanced expression states. Some SNPs were consistently associated with the unclassified group under both conditions, indicating their role as informative markers in this subgroup. Others showed specificity, being linked exclusively to either balanced or imbalanced expression, suggesting potential functional or regulatory roles. Similarly, several SNPs demonstrated ancestry-specific associations, particularly with the Black/African American group, reinforcing the interplay between germline variation, allele-specific expression, and population background in shaping transcriptional phenotypes.

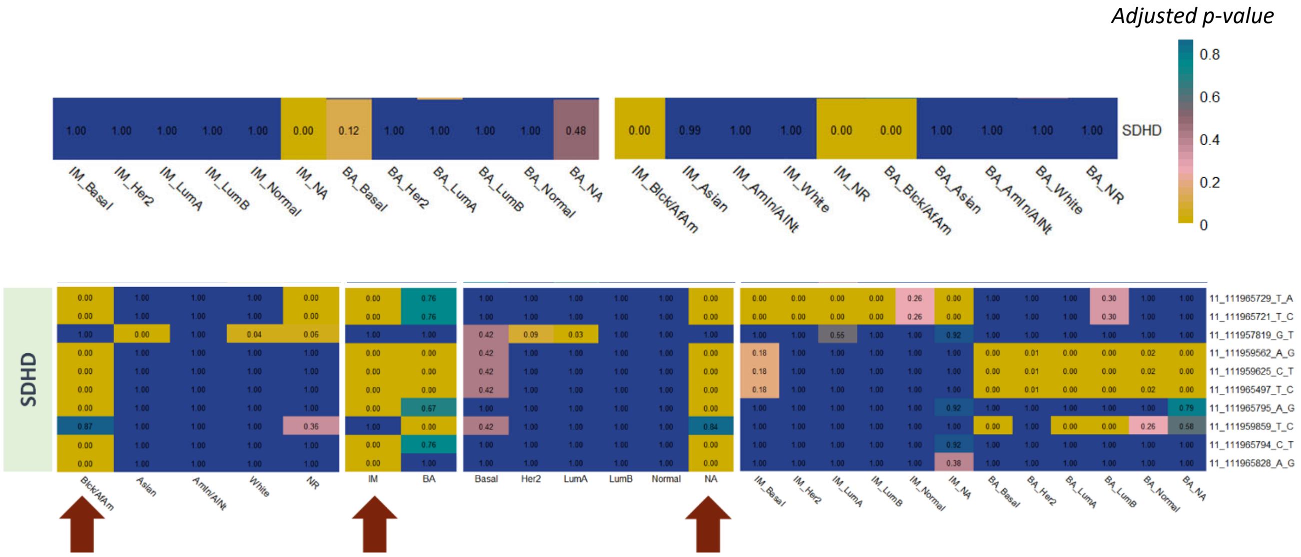

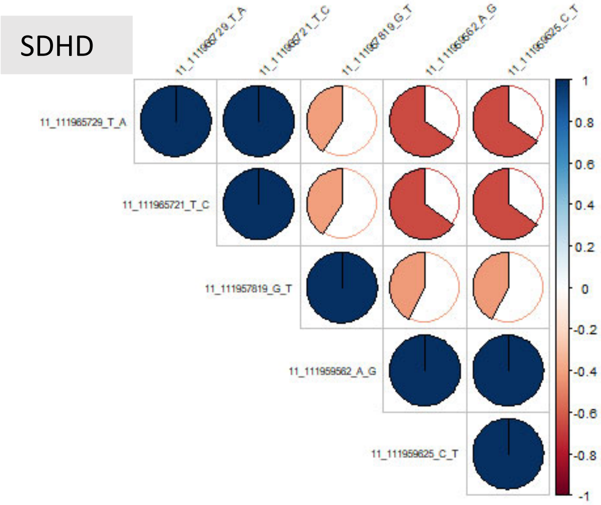


**Figure 14.** A) Association of balanced and imbalanced samples of SDHD gene with ethnicity and subtype. B) Association of phased SNIPs in SDHD with ethnicity and subtype; separate in balanced and imbalanced samples. C) The co-occurrence of SNIPs together.

##### *PAM 50 markers not classifying the NA group.*

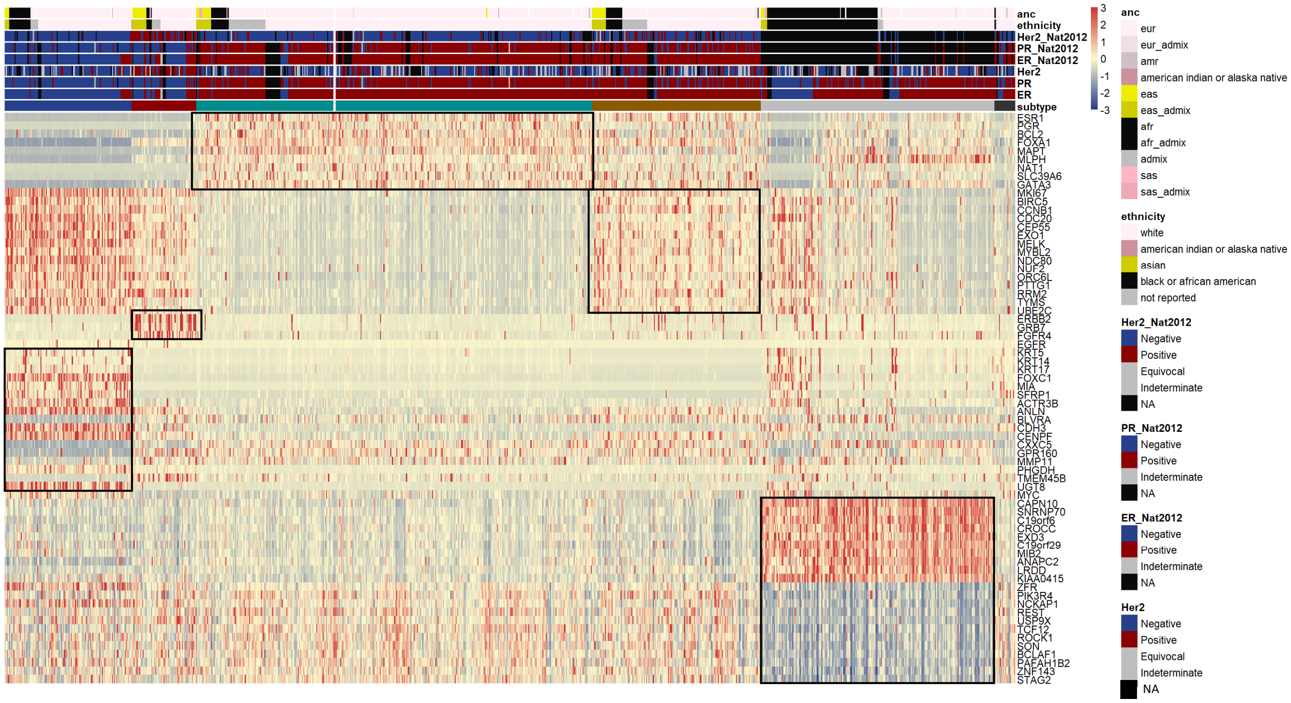


**Figure 15.** Expression profile of the marker genes of PAM50 + the potential marker genes identified for the unclassified tumors.

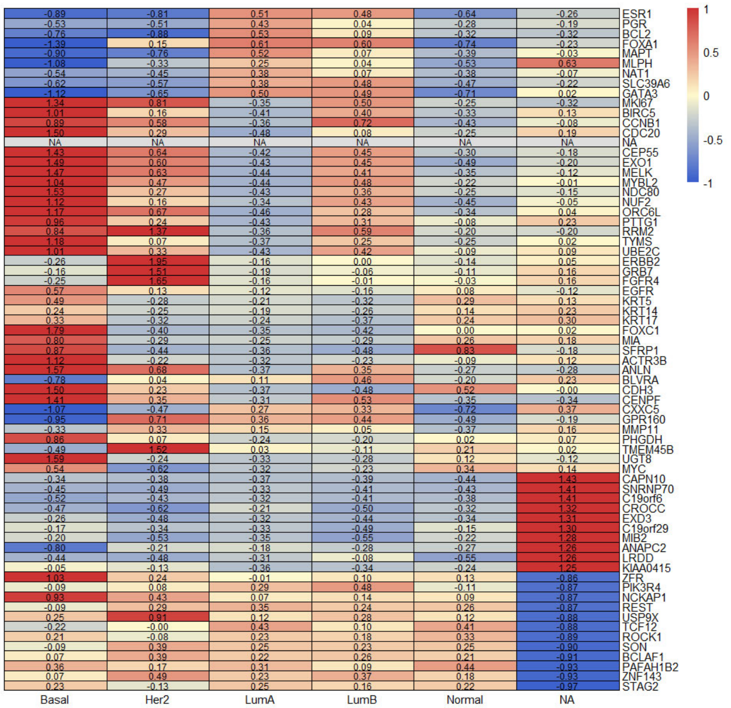


**Figure 16.** Mean z-score expression of the marker genes of PAM50 + the potential marker genes identified for the unclassified tumors.

#### ***TCGA with marker expressions.***

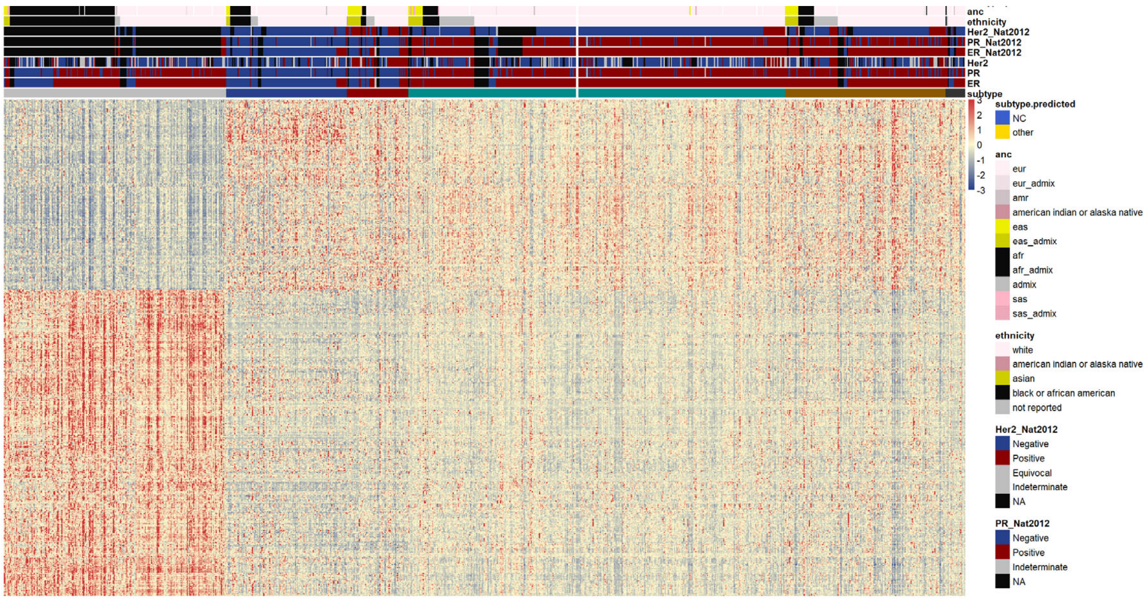


**Figure 17.** Expression profile of the potential marker genes of the unclassified tumors in TCGA; validated in METABRIC.
